# Comparison of endogenously expressed fluorescent protein fusions behaviour for protein quality control and cellular aging research

**DOI:** 10.1101/2021.01.25.428053

**Authors:** Kara L. Schneider, Adam Wollman, Thomas Nyström, Sviatlana Shashkova

## Abstract

The yeast Hsp104 protein disaggregase is often used as a reporter for misfolded or damaged protein aggregates and protein quality control and ageing research. Observing endogenously expressed Hsp104 fusions with fluorescent proteins is a popular approach to follow post stress protein aggregation, inclusion formation and disaggregation. Overall, such protein fusions used in molecular and microbiology research are often sparsely characterised. To address this issue, we performed a comparative assessment of Hsp104 fluorescent fusions function and behaviour. We provide experimental evidence that molecular behaviour may not only be altered by introducing a fluorescent protein tag but also varies depending on the fluorophore within the fusion. Although our findings are especially applicable to protein quality control and ageing research in yeast, similar effects and points may play a role in other eukaryotic systems.

## Introduction

Proper protein folding ensures maintenance of physiology and correct functioning of a cell and the entire organism [1]. Therefore, cells possess a protein quality control network to maintain protein homeostasis. This includes molecular chaperones that assist protein folding and prevent aggregation, degradation systems that recognise and remove terminally damaged proteins and disaggregases that, if necessary, ensure that misfolded proteins reacquire their native structure [2]. However, various factors such as environmental stress (heat shock, oxidative or UV/IR radiation) and ageing might cause disbalance in proteostasis followed by chronic expression of misfolded or aggregated proteins which is recognised as a hallmark of neurodegenerative (Alzheimer’s, Parkinson’s, Huntington’s diseases) and also some non-neurological disorders (Type 2 Diabetes, inherited cataract, some forms of atherosclerosis) [3]–[6].

In the budding yeast *Saccharomyces cerevisiae*, the protein disaggregase, Hsp104 heat-shock protein, binds to stress-induced protein aggregates to disassemble and reactivate them. It has been suggested that Hsp104-bound aggregates that appear immediately in response to a stress, over time, coalesce into specific protein inclusions, such as IPOD (Insoluble-Protein-Deposit), INQ (Intra-Nuclear-Quality-Control) and JUNQ (Juxta-Nuclear-Quality-Control) [7]–[9]. Similarly, age-induced protein aggregates are also targeted by Hsp104 [10]. Thus, Hsp104 is a commonly used reporter for misfolded or damaged protein aggregates that can be monitored under the microscope as intracellular foci.

The discovery of the green fluorescent protein, GFP, from the jellyfish *Aequorea victoria* [11], revolutionised experimental approaches in molecular biology. Since the first reports of the gene for GFP had been cloned and sequenced [12], it became widely used as a reporter for protein studies. A large number of mutations have now been introduced into the wild-type GFP, as well as the creation of newly engineered fluorescent proteins in order to improve biophysical characteristics as well as obtain new colours [13]. However, while fluorescent proteins allow direct visualisation of proteins of interest, numerous technical limitations still remain [13], [14]. For instance, experiments that require long-term illumination for longer observations of studied processes are significantly restricted by maturation times and photostability of fluorescent protein. Moreover, such tags significantly increase the overall size of the protein construct, which might affect its natural molecular conformations, hence, its behaviour and function. Therefore, the choice of the most suitable fluorescent tag largely depends on the type of the experiment as well as information one wants to obtain.

Here, we performed a comparative assessment of endogenously expressed Hsp104-fluorescent protein fusions under control of the native *HSP104* promoter. We report comparisons of a version of an enhanced GFP used in the GFP-tagged protein library (GFP) [15], a monomeric form of GFP (mGFP) and two recently described bright monomeric fluorescent proteins: green mNeonGreen (derived from a tetrameric fluorescent protein from cephanolochordate *Branchiostoma lanceolatum*) [16] and red mScarlet-I [17]. We provide comprehensive data that can be utilised for choosing a fluorescent protein for *in vivo* and *in vitro* experiments on the budding yeast in protein quality control and ageing research.

## Materials and Methods

### Media and growth conditions

Cells from frozen stocks were pre-grown on standard YPD medium plates (20 g/L Bacto Peptone, 10 g/L Yeast Extract) supplemented with 2% glucose (w/v) at 30°C. For liquid cultures, cells were grown in Yeast Nitrogen Base (YNB) medium (1x Difco™ YNB base, 1x Formedium™ Complete amino acid Supplement Mixture, 5.0 g/L ammonium sulfate, pH 5.8-6.0) supplemented with 2% glucose (w/v), at 30°C, 180 rpm.

### Strain construction

We created a number of novel yeast strains expressing fluorescent Hsp104 by introducing *mGFP-HIS3*, *mNeonGreen-HIS3* and *mScarlet-I-LEU2* fragments flanked on their 5’- and 3’-ends with ~50 bp sequences up- and downstream of the Hsp104 *STOP* codon, respectively. The *mGFP-HIS3* fragment was amplified from the pmGFP-S plasmid [18]. pmNG-S and pmScI-S plasmids were created for this study by introducing mNeonGreen and mScarlet-I sequences into YDp-H and YDp-L vectors, respectively. PCR reaction mixes with amplified fragments were introduced into the yeast genome by standard LiAc protocol [19]. Successful transformants were verified by confirmation PCR and standard fluorescence microscopy. Full list of strains and plasmids used in this study is presented on Table 1 and Table 2, respectively.

**Table 1.**

| Name | Genotype | Source/reference |
| --- | --- | --- |
| BY4741 wild type | *MATa his3Δ1 leu2Δ0 met15Δ0 ura3Δ* | Nyström collection |
| *hsp104Δ* | *MATa hsp104Δ::kanMX4 his3Δ1 leu2Δ0 met15Δ0 ura3Δ0* | Nyström collection |
| Hsp104-GFP | *MAT a his3Δ1 leu2Δ0 met15Δ0 ura3Δ0 HSP104-GFP:HIS3MX6* | [15] |
| Hsp104-mGFP | *HSP104-GFPmut3 (S65G, S72A, A206K)-HIS3* (in BY4741) | this study |
| Hsp104-mNG | *HSP104-mNeonGreen-HIS3* (in BY4741) | this study |
| Hsp104-mScl | *HSP104-mScarlet-I-LEU2* (in BY4741) | this study |
| Tom70-GFP | *MAT a his3Δ1 leu2Δ0 met15Δ0 ura3Δ0 TOM70-GFP:HIS3MX6* | [15] |
| Tom70-mGFP | *TOM70-GFPmut3 (S65G, S72A, A206K)-HIS3* (in BY4741) | this study |
| Tom70-mNG | *TOM70-mNeonGreen-HIS3* (in BY4741) | this study |
| Tom70-mScl | *TOM70-mScarlet-I-LEU2* (in BY4741) | this study |

**Table 2.**

| Name | Description | Source/Reference |
| --- | --- | --- |
| pmGFP-S | *GFP S65G, S72A, A206K in YDp-H* | [18] |
| pmNG-S | *mNG in YDp-H* | this study |
| pmSc-S | *mScl in YDp-L* | this study |
| YDp-H | *HIS3* | [20] |
| YDp-L | *LEU2* | [20] |

### Growth curves

For the growth curve experiments, strains were pre-grown overnight in YNB medium supplemented with 2% glucose. Cells were then sub-cultured to OD_600_ ~0.01 and placed onto a 24-well plate for monitoring the growth. Absorbance measurements at 600 nm were taken every 60 min for more than 50 -72 h. Doubling times and the length of the Lag phase were estimated using the PRECOG software [21].

### Mature fraction of fluorescent proteins

Yeast strains were pre-grown overnight in YNB complete medium supplemented with 2% glucose. Cells were then sub-cultured to OD_600_~0.2 and grown for 4h (until mid-logarithmic phase). To inhibit protein synthesis, the cultures were subjected to 100μg/ml cycloheximide (Sigma Aldrich) for 1h at room temperature, protected from light). The cells were placed onto a 1% agarose pad prepared using Geneframes (Thermo Scientific) supplemented with 1x CSM, 1x YNB, 2% glucose and 250 μg/ml cycloheximide. Images were acquired at room temperature. Mature fraction of fluorophores within protein fusions was estimated as described previously [22].

### Aggregate clearance

Cells pre-grown to OD_600_ ~0.4-0.5 were subjected to 42°C heat shock for 30 min. For recovery, cells were then placed into 30°C, 180 rpm. To follow aggregates formation and clearance, samples were taken prior to the heat shock, immediately after, and after 30, 60 and 90 min of recovery. Cells were fixed in 3.7% formaldehyde (Scharlau) and imaged in z-stacks. The number of aggregates per cell was scored manually using open ImageJ FiJi 2.1.0/1.53c software, Сell Сounter Plugin. Efficiency of aggregate clearance was calculated for each time point after the heat shock as a percentage of cells without any fluorescent foci.

### Western blotting

Cells were pre-grown overnight, sub-cultured to OD_600_ = 0.1 and grown to OD_600_ ~ 0.45-0.55. The cells were then heat shocked at 42°C for 30 min and sampled by collecting 50 ml culture before heat shock, right after and then 60 and 90 min after recovery at 30°C. Proteins were extracted by boiling the cells in Laemmli buffer [23], [24]. Protein concentration was determined using Pierce 660 nm assay (Thermo Scientific). 20 μg of proteins were loaded per lane on a 10% Criterion TGX Precast Gel (Bio-Rad Laboratories) and resolved in TGX buffer (Bio-Rad) at 60 V for ~1h and then at 130-150 V until the stain line ran out of the gel. Blotting was performed with the Trans Blot Turbo Transfer System onto a 0.2 μm Nitrocellulose membrane (Bio-Rad). The membrane was then blocked using Intercept (PBS) blocking buffer (LI-COR Biosciences) for 1 h followed by probing with primary anti-Hsp104 (ab69549, Abcam, 1:2000 dilution) and anti-Pgk1 (ab90787, Abcam, 1:20000 dilution) antibodies for overnight at 4°C. The membrane was then washed in PBS with 0.1% Tween-20 and incubated with goat anti-rabbit IRDye 800CW and goat anti-mouse IRDye 680RD (LI-COR; 1:20000 dilution) secondary antibodies for 1 h. After washing, the membrane was scanned using the LI-COR Odyssey Infrared scanner using 800 and 700 nm channels, respectively. Hsp104 protein expression levels were calculated based on the background corrected band intensity measured by ImageJ FiJi 2.1.0/1.53c software normalised to the control Pgk1 protein. Protein expression change was calculated as a change relative to the initial (before the heat stress) Hsp104 protein levels.

### Heat tolerance assay

Overnight pre-grown strains were sub-cultured and grown to OD600 ~0.5. The strains were sampled for spot test as a 30°C control. Remaining cultures were split into two groups and heat-stressed in a 37°C water bath. One of the sample groups was collected after 30 min as a 37°C sample, the other group was further heat-treated in a 50°C water bath for 30 min. All strains were placed onto YPD plates in 5 10-fold dilution 5 μl spots and allowed to grow at 30°C for 3 days.

### Epifluorescence microscopy

To determine Hsp104 localisation, strains were grown in YNB complete medium supplemented with 2% glucose (weight per volume) overnight until the stationary phase. Live cells were then stained with 4’,6-diamidino-2-phenylindole (DAPI) (final concentration 10 μg/ml) for 5 min and directly visualised using a Zeiss Axio Observer.Z1 inverted microscope with Apotome and Axiocam 506 345 camera, and a Plan-Apochromat 100x/1.40 Oil DIC M27 objective.

For heat stress recovery experiments, every field of view was visualised as a Z-stack of 10 images through the range of 7 μm.

For timelapse microscopy, the strain expressing Hsp104-mSc-I was grown in YNB complete medium supplemented with 2% glucose, sub-cultured to OD_600_ ~0.45 and subjected to 30 min heat-shock at 42°C. The cells were then gently spun down and placed onto a 1% agarose pad perfused with YNB medium supplemented with 2% glucose (w/v) and sealed with a coverslip as described previously [24]. The sample was imaged in 11 Z-stacks every 5 min for 90 min at 30°C using the TempModule S1 (Zeiss), Y-module S1 (Zeiss), Temperable insert S1 (Zeiss), Temperable objective ring S1 (Zeiss), and Incubator S1 230V (Zeiss) to maintain the temperature. The focus was set manually and maintained using the software autofocus with SR101 (593 nm) as a reference channel. The timelapse images were processed with Fiji software using ImageJ plugin BleachCorrect V.2.0.2 with simple ratio and the manual drift correction plugin [25]. For representation, enhancements in brightness/contrast were used.

## Results

### Fluorescent tags do not affect growth characteristics

Fluorescent tags have been reported to affect yeast growth fitness via altering protein function and intracellular localisation [26]. We first examined weather fluorescent labelling of Hsp104 had any impact on growth rates of yeast cultures. We compared the BY4741 wild type and four strains with endogenously labelled Hsp104: Hsp104-GFP, Hsp104-mGFP, Hsp104-mNeonGreen and Hsp104-mScarlet-I. The *hsp104Δ* strain was also used as a reference to Hsp104 dysfunction. The cultures were monitored for 72 h and the growth was measured based on the absorbance at 600 nm. The growth curve profile was identical for all strains tested (Figure 1A). No significant difference (Student’s *t-test*) between doubling times was observed between all cultures (Figure 1B). We estimated the length of the lag phase to be similar (Student’s *t-test*) and slightly over 4 h across all strains tested (Figure 1B).

**Figure 1.**
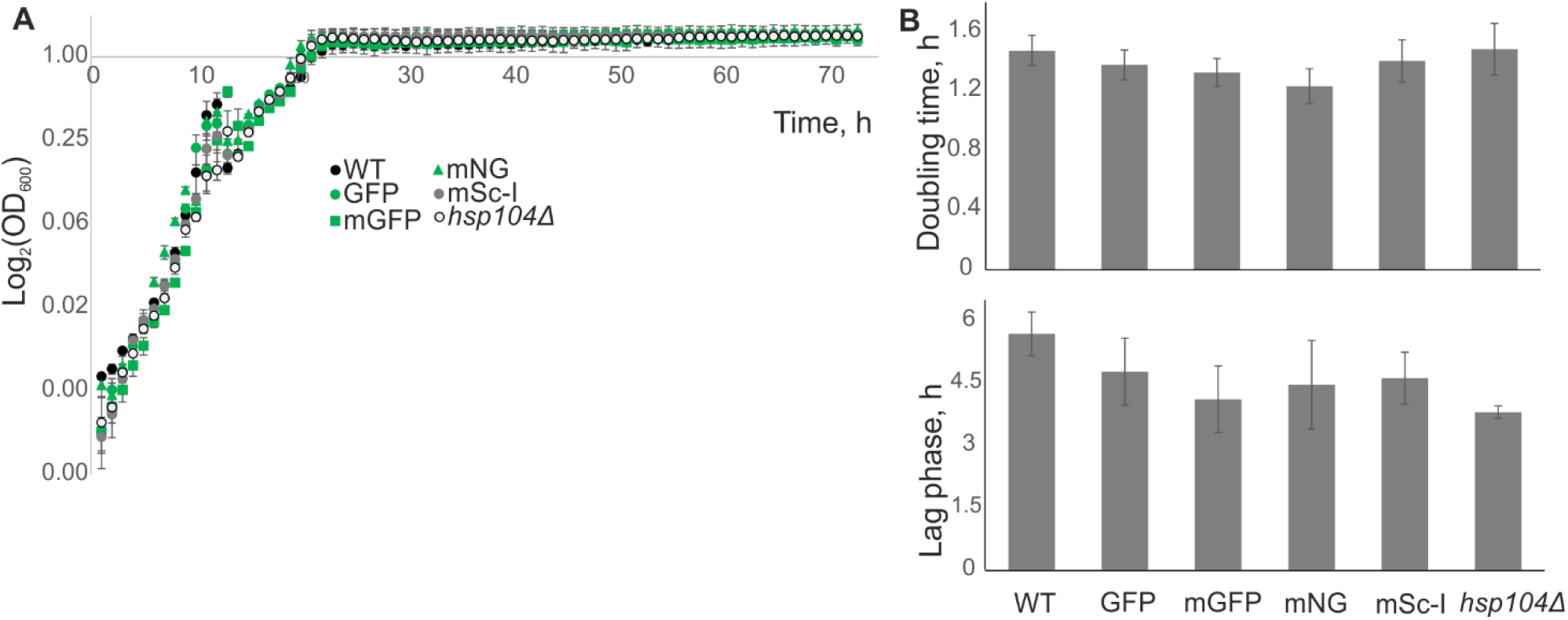
Growth characteristics of the BY4741 wild type (WT), cells expressing endogenous constructs Hsp104-GFP (GFP), Hsp104-mGFP (mGFP), Hsp104-mNeonGreen (mNG), Hsp104-mScarlet-I (mSc-I) and a deletion *hsp104Δ* strain. **A.** Growth curves of yeast cultures. Each time point is a mean value of technical replicates of one strain. Error bars represent standard error of mean. **B.** (top) Mean doubling time of different strains, (bottom) average length of the lag phase for different strains. Standard error of mean error bars. Data from one of three representative experiments is shown.

### Fluorescent tags affect intracellular localisation of Hsp104

According to the Yeast GFP Fusion Localization Database, under standard conditions, the Hsp104 protein is located in the cytoplasm. However, several studies report both nuclear and cytoplasmic localisation of Hsp104 in unstressed cells [27]. We performed live cell imaging of cell in stationary phase to determine whether the subcellular localisation of Hsp104 differs depending on the fluorescent tag. While Hsp104-GFP and Hsp104-mSc-I were solely cytoplasmic, Hsp104-mGFP and Hsp104-mNG exhibited also nuclear localisation (Figure 2).

**Figure 2.**
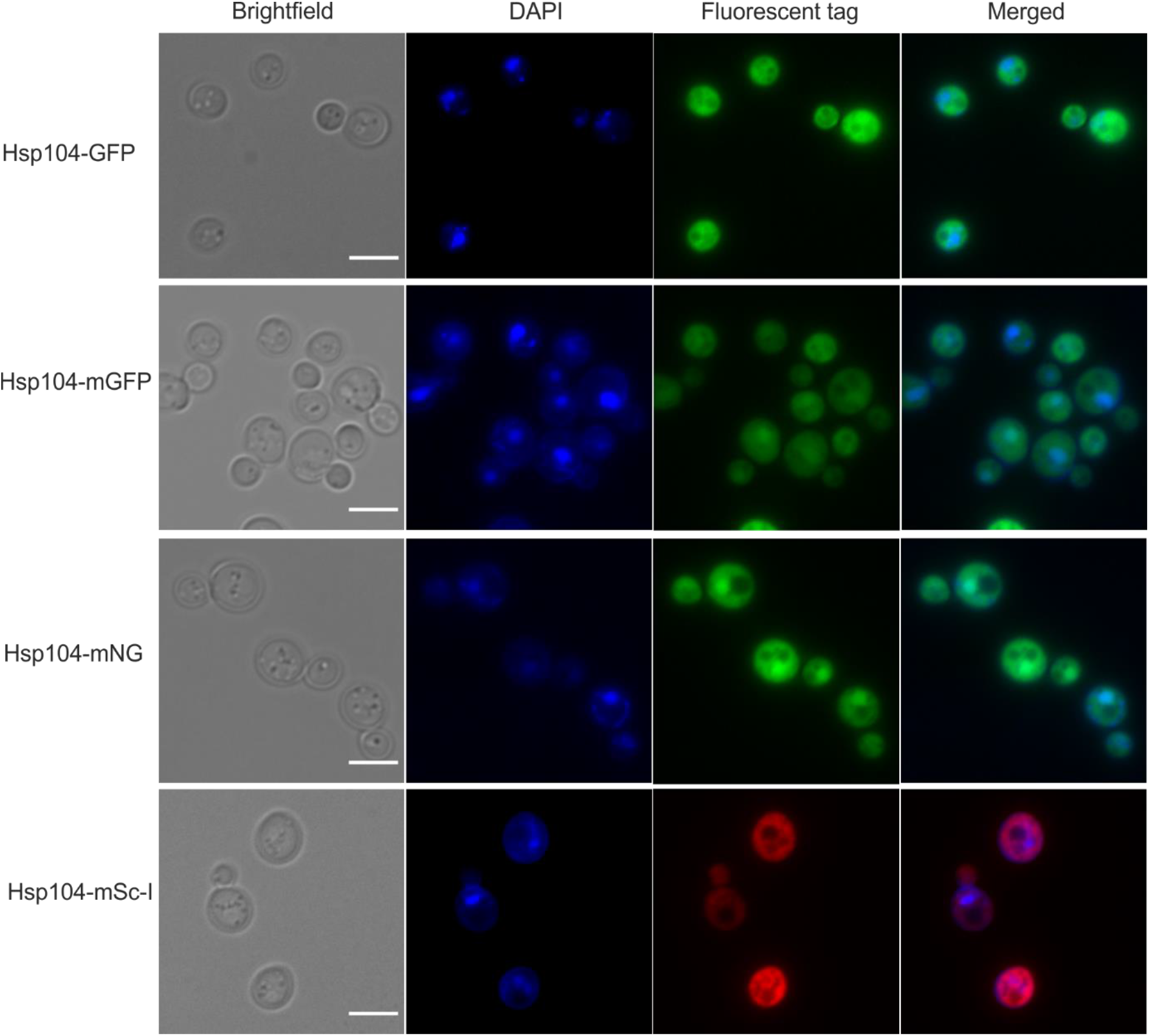
Subcellular localisation of Hsp104 tagged with fluorophores. Live images of overnight grown cells stained with DAPI for nuclear detection. Scale bar 5μm.

While the GFP-labelled strain originates from a previously created library [15], other fluorescent strains were created by introducing the fluorescent protein encoding sequence into the genome via homologous recombination. To obtain the insertion fragments, we amplified the DNA fragments by the polymerase chain reaction (PCR), a method which may provide errors in the final product due to thermal damage of DNA or editing errors during DNA copying [28]. Such mutations may cause changes in protein sequence, hence affect its behaviour, which could result in altered localisation. To explore this possibility, we sequenced DNA of the wild type and newly designed fluorescent fusions of Hsp104. Sequencing results did not indicate any PCR-induced mutations within cells. Therefore, subcellular distribution of Hsp104 fusions seems to be defined by fluorophores.

### Efficiency of aggregate clearance

The Hsp104 chaperone is widely used in ageing and proteopathy research as a marker of misfolded or damaged protein aggregates. In yeast, *S. cerevisiae*, Hsp104 binds to stress-induced misfolded proteins and assists their refolding. To test whether the rate of protein aggregate removal differs and depends on the fluorescent tag, we performed a clearance assay of heat-induced aggregates.

After 30 min at 42°C, all the cells showed clear response with high numbers of fluorescent foci corresponding to heat stress-induced damaged protein aggregates. While all strains showed significant decrease in the amounts of fluorescent spots 60 min after the cells were brought to their standard growth conditions (30°C, 180 rpm), only Hsp104-mSc-I strain exhibited almost 100% clearance as opposed to 30-40% for other strains (Figure 3A). Interestingly, while other strains reached complete aggregate removal at 90 min recovery, by this point the number of cleared cells expressing Hsp104-mSc-I significantly decreases (Student’s *t-test*, p<0.05). More detailed analysis revealed that the main source of this artefact was mother cells while the daughter cells stayed aggregates free (Figure 3B). Like other proliferating cells, yeast undergo asymmetric cell division which allows for asymmetric segregation of damaged proteins meaning the mother cell retains protein aggregates, and a new daughter cell is produced free of damage and with a full replicative potential [2]. Thus, one possible mechanism of how daughter cells are cleared from aggregated proteins is by dragging them back into the mother cell. To further investigate the Hsp104-mSc-I behaviour during the recovery, we performed timelapse microscopy to follow the fluorescent foci within the mother and the daughter cells after 30 min exposure to the 42°C heat stress (Movie 1). While we can observe mother cells retrieving the damaged protein aggregates back from the daughter cells (Figure 3C, Movie 1), the mSc-I foci reappearing in the mothers by 90 min recovery, seem to be originated within the mother cells (Figure 3D). The overall pattern of aggregate removal from the daughter cells was similar between all fluorescent strains tested (Figure 3E) with complete clearance of cells expressing Hsp104-mSc-I already after 60 min recovery, while the rest reached this level 30 min later. Interestingly, a small percent of daughter cells expressing mGFP appeared to be aggregate free already immediately after 30 min exposure to the heat stress.

**Figure 3.**
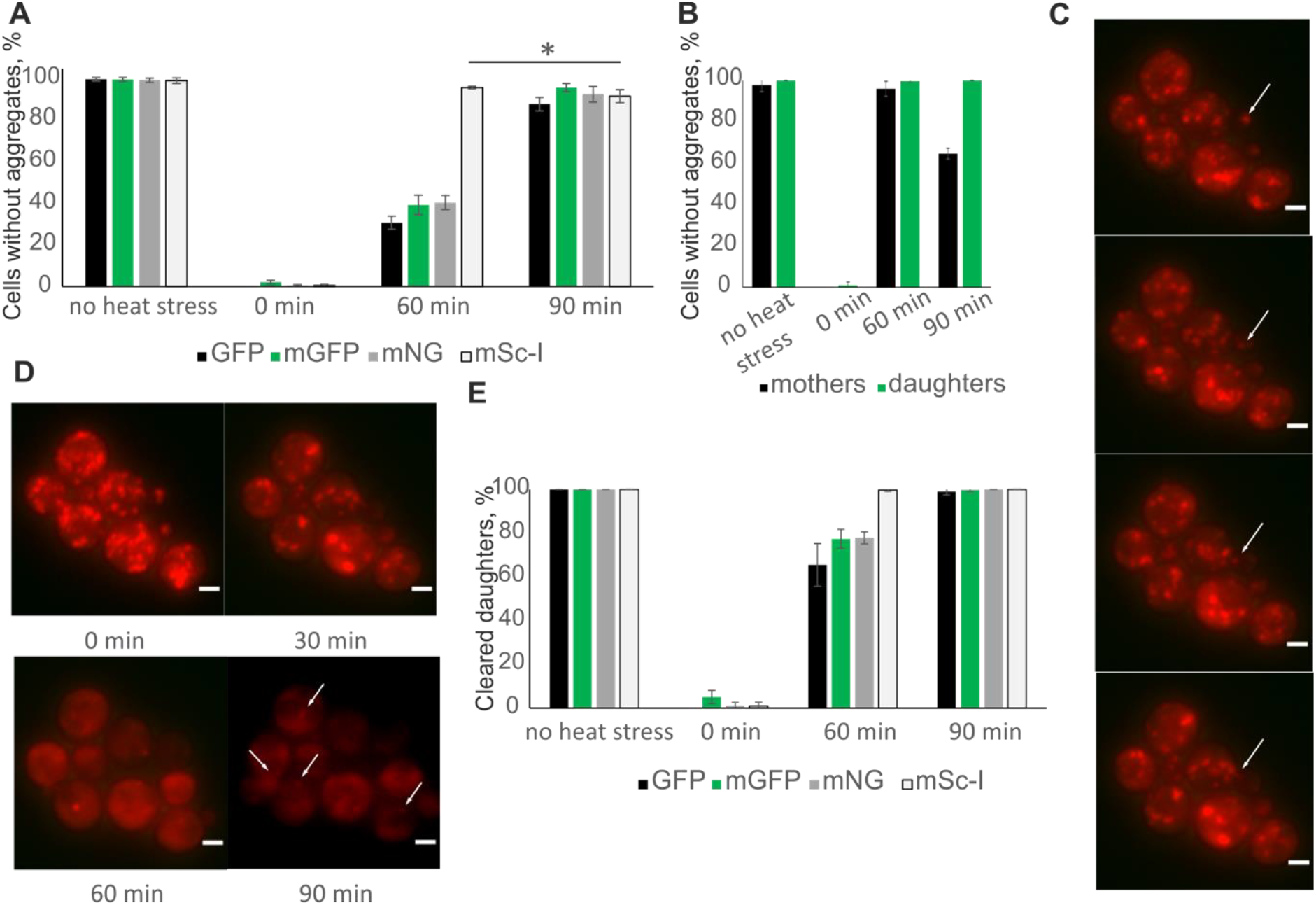
Heat-induced damaged protein aggregates removal efficiency. **A**. Mean percent of cells without aggregates before the stress, immediately after, 60 and 90 min after cells were returned into their standard growth conditions (recovery). Cells expressing Hsp104 fused with GFP (black), mGFP (green), mNeonGreen (grey) and mScarlet-I (white) were tested. Error bars represent standard deviation. * p<0.05, Student’s t-test. **B**. Mean percent of mother (black) and daughter (green) cells expressing Hsp104-mScarlet-I before stress, immediately after, 60 and 90 min recovery. Standard deviation error bars. **C**. Representative images of cells with Hsp104-mSc-I recovering after the heat stress (15-30 min). White arrows indicate a fluorescent focus being dragged from the daughter to the mother cell. Scale bar 2 μm. **D**. Representative image of Hsp104-mSc-I cells recovering from the heat stress (0-90 min). White arrows indicate fluorescent foci reappearing in the mother cells after 90 min recovery. Scale bar 2 μm **E**. Mean percent of daughter cells without aggregates before and after heat stress. Cells expressing Hsp104 fused with GFP (black), mGFP (green), mNeonGreen (grey) and mScarlet-I (white) were tested. Standard deviation error bars.

### Hsp104 and heat tolerance

To test whether the differences in heat-induced foci formation is due to the fact that a fluorescent tag alters protein expression, we examined Hsp104 levels in cells subjected to the heat stress followed by recovery at the standard conditions (30°C, 180 rpm). Our data show a drastic increase of the Hsp104 expression upon exposure to the heat stress (Figure 4A) which is consistent with our expectations based on previous reports [29], [30]. Fluorescently tagged Hsp104 exhibited lower initial levels of protein expression (Figure 4A). While this could be an effect of potentially lower antibody binding efficiency to Hsp104 fusions, the Hsp104 protein expression change in response to the high temperature was smaller for fluorescent Hsp104 compared to that of the unlabelled Hsp104 in the wild type strain (Figure 4B).

**Figure 4.**
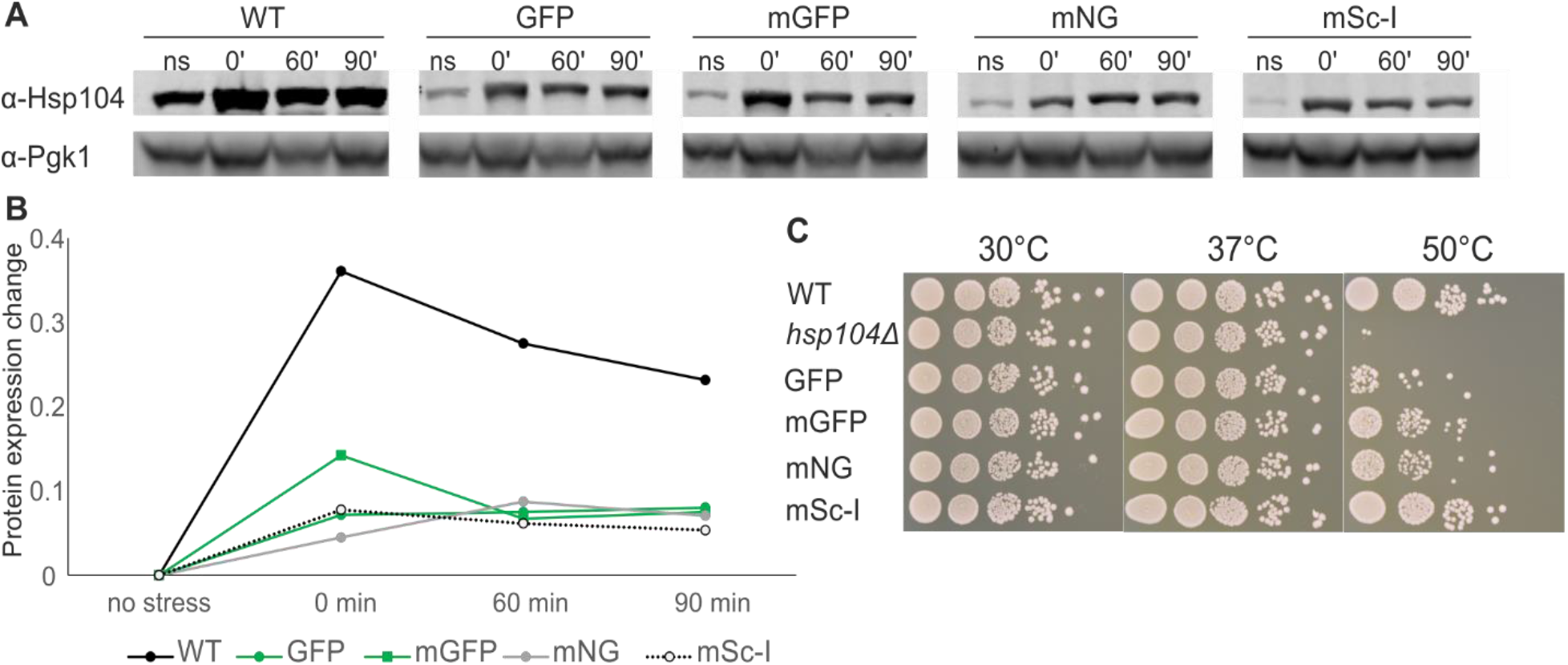
**A**. Hsp104 protein expression levels from total protein extracts before (ns) and right after (0’) to the heat stress as well as 60 and 90 min after recovery at 30°C, 180 rpm, in the wild type (WT) strain and cells expressing Hsp104 tagged with GFP, mGFP, mNeonGreen (mNG) and mScarlet-I (mSc-I). **B**. Hsp104 protein expression change right after the heat stress, and 60 and 90 min recovery relative to the initial expression level in the wild type (black circles) and cells with Hsp104 labelled with GFP (green circles), mGFP (green squares), mNG (grey circles) and mSc-I (empty circles). Quantification of the western blot presented on panel A. One representative experiment of three biological replicates. **C**. 3 days recovery at 30°C of the wild type (WT), *hps104Δ* and strains expressing Hsp104 fused with GFP, mGFP, mNG and mSc-I after the thermal insult at 50°C for 30 min with a pretreatment at 37°C for 30 min.

Overall, fluorescent fusions did not disrupt the Hsp104 function in cell survival after the heat insult (Figure 4C). As a control for the total loss of function, we used the *hsp104Δ* strain which failed to survive at high temperatures even with a prior treatment at 37°C. Such pretreatment has been suggested to be essential for cellular recovery after the heat insult. Interestingly, while recovery rates for strains with green fluorophores were lower compared to the unlabelled strain, cells carrying the Hsp104-mSc-I fusion survived as well as the wild type.

## Discussion

With the development of fluorescent microscopy, the use of fluorophores as protein fusions or individual particles within cells became an invaluable tool in modern cell and molecular biology research. While allowing for direct visualisation of proteins of interest directly inside the cell, fluorescent tags increase the overall protein size and might alter its function through, for example, disrupting native protein conformation or changing accessibility of essential domains and regions to other molecules [13]. Despite being widely used, very few studies examined biophysical characteristics of fluorescent proteins within living systems [31]–[33]. Fluorescent proteins within protein fusions are even more sparsely characterised. Here we show that the behaviour of the protein of interest cannot only be altered by a tag but also varies depending on a fluorophore.

While we did not observe any effect on cell cultures growth characteristics, we report alterations in Hsp104 subcellular localisation. Several C-terminal point mutations from lysine to alanine that inhibit nuclear localisation of Hsp104 have been suggested [27]. Experimental and computational studies on amino acid substitutions indicate their drastic effects on protein folding, stability and protein-protein interactions [34]. However, we did not identify any previously reported *HSP104* mutations. Therefore, fluorescent tags themselves may alter protein conformation which may affect behaviour and localisation. While we show that proteins behaviour and localisation changes depending on a fluorescent label, a detailed structural analysis is beyond the scope of this study, but it is required to provide a definitive answer on how the conformation of a protein of interest is modified within fluorescent fusions *in vivo*.

In yeast cultures, cellular rejuvenation and a life span reset is achieved during cellular division when the mother cells maintain all misfolded and damaged proteins, and a new daughter cell is created free of damage. However, to deal with stress-induced damage, cells possess a cohesive protein quality control (PQC) system, where the Hsp104 protein disaggregase assists damaged protein aggregates clearance via their disassembly and protein refolding [35], [36]. Such protein refolding and reactivation is essential for longevity of living organisms, thus, overexpression of the yeast Hsp104 has been shown to prolong the lifespan in mice models [37]. Our data clearly indicate heat-induced aggregate clearance within 90 min after the exposure to the high temperature. Interestingly, we could observe Hsp104-mGFP cells free from aggregates, mainly daughter cells, immediately after the stress conditions were removed. This corresponds to higher level of the Hsp104 protein expression in this strain at this time point compared to other strains carrying fluorescently labelled Hsp104. Having similar protein expression levels, only Hsp104-mSc-I construct showed complete aggregate clearance already one-hour post stress. Whether this fusion possesses increased Hsp104 activity or it triggers other components that protect young cells from damage, remains to be investigated. Surprisingly, 90 min post stress we observe an increasing number of mother cells (ca 30%) with one aggregate. While we did capture protein aggregates being dragged from a daughter cell at earlier times after the heat stress, those newly appeared fluorescent foci have their origin within the mother cells. Prior to our experiments, we estimated a dark immature fraction of fluorescent proteins within endogenously expressed fluorescent fusions. To overcome low Hsp104 expression level at the normal conditions, we created and analysed the strains also expressing fluorescent fusions with Tom70, a mitochondrial protein. While all expressed GFP, mGFP and mNeonGreen proteins were fully matured, we estimated 10% of mScarlet-I dark fraction, and its maturation time about 30 min which is consistent with previously reported data and is longer than that of GFP, mGFP and mNeonGreen [38]. Therefore, newly appearing foci are likely to be age-related, hence while enhancing damage clearance efficiency in daughter cells, the Hsp104-mSc-I construct provides more stress within the mother cell. Fast daughter cells clearance seems to be the reason of a better and comparable to the wild type heat insult tolerance of Hsp104-mSc-I compared to other fluorescent strains tested.

This work highlights the importance of characterising the effects that fluorescent tags can have on protein function, specifically for the field of proteostasis and ageing, which largely relies on fluorescence microscopy. Our data clearly indicate that the behaviour and function of the protein of interest can be severely affected by fluorescent labels. We discuss the issues that researchers may consider upon choosing fluorescent proteins for their experimental approach. In addition to monitoring endogenous proteins using the general disaggregase Hsp104, many misfolding reporters are available to be introduced into the cell. These are known to be affected by fluorescent tagging as well, which should be taken into account when selecting tools to study protein quality control [36]. Although our findings are especially applicable to protein quality control and ageing research in the budding yeast *S. cerevisiae*, similar effects and points may play a role in other eukaryotic systems.

## Funding sources

This work has been supported by the Marie Curie Alumni Association, the Royal Society Newton International Fellowship Alumni (AL\191025), Knut and Alice Wallenberg Foundation (KAW 2017-0091, KAW 2015.0272), Swedish Research Council (VR 2019-03937).

## Notes

### Competing Interest Statement

The authors have declared no competing interest.

## References

[1] C. M. Dobson, “Principles of protein folding, misfolding and aggregation,” Semin Cell Dev Biol, vol. 15, no. 1, pp. 3–16, 2004.

[2] S. M. Hill, S. Hanzén, and T. Nyström, “Restricted access: spatial sequestration of damaged proteins during stress and aging,” EMBO Rep., vol. 18, no. 3, pp. 377–391, Mar. 2017.

[3] C. López-Otín, M. A. Blasco, L. Partridge, M. Serrano, and G. Kroemer, “The hallmarks of aging,” Cell, vol. 153, no. 6. Cell Press, p. 1194, 06-Jun-2013.

[4] R. A. Brewer, V. K. Gibbs, D. L. Smith, and Jr., “Targeting glucose metabolism for healthy aging,” Nutr. Heal. Aging, vol. 4, no. 1, p. 31, 2016.

[5] F. Ursini, K. J. A. Davies, M. Maiorino, T. Parasassi, and A. Sevanian, “Atherosclerosis: Another protein misfolding disease?,” Trends in Molecular Medicine, vol. 8, no. 8. Elsevier, pp. 370–374, 01-Aug-2002.

[6] K. L. Moreau and J. A. King, “Protein misfolding and aggregation in cataract disease and prospects for prevention,” Trends in Molecular Medicine, vol. 18, no. 5. NIH Public Access, pp. 273–282, May-2012.

[7] D. Kaganovich, R. Kopito, and J. Frydman, “Misfolded proteins partition between two distinct quality control compartments,” Nature, vol. 454, no. 7208, pp. 1088–1095, 2008.

[8] S. Escusa-Toret, W. I. M. Vonk, and J. Frydman, “Spatial sequestration of misfolded proteins by a dynamic chaperone pathway enhances cellular fitness during stress,” Nat. Cell Biol., vol. 15, no. 10, pp. 1231–1243, Oct. 2013.

[9] S. B. Miller et al., “Compartment‐specific aggregases direct distinct nuclear and cytoplasmic aggregate deposition,” EMBO J., vol. 34, no. 6, pp. 778–797, Mar. 2015.

[10] N. Erjavec, L. Larsson, J. Grantham, and T. Nyström, “Accelerated aging and failure to segregate damaged proteins in Sir2 mutants can be suppressed by overproducing the protein aggregation-remodeling factor Hsp104p,” Genes Dev, vol. 21, no. 19, pp. 2410–2421, 2007.

[11] O. Shimomura, F. H. Johnson, and Y. Saiga, “Extraction, Purification and Properties of Aequorin, a Bioluminescent Protein from the Luminous Hydromedusan, Aequorea,” J. Cell. Comp. Physiol., vol. 59, no. 3, pp. 223–239, Jun. 1962.

[12] D. C. Prasher, V. K. Eckenrode, W. W. Ward, F. G. Prendergast, and M. J. Cormier, “Primary structure of the Aequorea victoria green-fluorescent protein,” Gene, vol. 111, no. 2, pp. 229–233, Feb. 1992.

[13] S. Shashkova and M. C. Leake, “Single-molecule fluorescence microscopy review: shedding new light on old problems,” Biosci. Rep., vol. 37, no. 4, p. BSR20170031, Aug. 2017.

[14] S. Shashkova and M. C. Leake, “Systems biophysics: Single-molecule optical proteomics in single living cells,” Curr. Opin. Syst. Biol., vol. 7, 2018.

[15] W. K. Huh et al., “Global analysis of protein localization in budding yeast,” Nature, vol. 425, no. 6959, pp. 686–691, Oct. 2003.

[16] N. C. Shaner et al., “A bright monomeric green fluorescent protein derived from Branchiostoma lanceolatum.,” Nat. Methods, vol. 10, no. 5, pp. 407–9, May 2013.

[17] D. S. Bindels et al., “MScarlet: A bright monomeric red fluorescent protein for cellular imaging,” Nat. Methods, vol. 14, no. 1, pp. 53–56, Dec. 2016.

[18] A. J. M. Wollman, S. Shashkova, E. G. Hedlund, R. Friemann, S. Hohmann, and M. C. Leake, “Transcription factor clusters regulate genes in eukaryotic cells,” Elife, vol. 6, p. e27451, Aug. 2017.

[19] R. D. Gietz and R. H. Schiestl, “Frozen competent yeast cells that can be transformed with high efficiency using the LiAc/SS carrier DNA/PEG method.,” Nat. Protoc., vol. 2, no. 1, pp. 1–4, Jan. 2007.

[20] G. Berben, J. Dumont, V. Gilliquet, P.-A. A. Bolle, and F. Hilger, “The YDp plasmids: A uniform set of vectors bearing versatile gene disruption cassettes forSaccharomyces cerevisiae,” Yeast, vol. 7, no. 5, pp. 475–477, Jul. 1991.

[21] L. Fernandez-Ricaud, O. Kourtchenko, M. Zackrisson, J. Warringer, and A. Blomberg, “PRECOG: A tool for automated extraction and visualization of fitness components in microbial growth phenomics,” BMC Bioinformatics, vol. 17, no. 1, p. 249, Jun. 2016.

[22] S. Shashkova, A. Wollman, S. Hohmann, and M. C. Leake, “Characterising Maturation of GFP and mCherry of Genomically Integrated Fusions in Saccharomyces cerevisiae,” BIO-PROTOCOL, vol. 8, no. 2, p. e2710, Jan. 2018.

[23] U. K. Laemmli, “Cleavage of structural proteins during the assembly of the head of bacteriophage T4,” Nature, vol. 227, no. 5259, pp. 680–685, 1970.

[24] S. Shashkova, M. Andersson, S. Hohmann, and M. C. Leake, “Correlating single-molecule characteristics of the yeast aquaglyceroporin Fps1 with environmental perturbations directly in living cells,” Methods, May 2020.

[25] K. Miura, C. Rueden, M. Hiner, J. Schindelin, and J. Rietdorf, “ImageJ Plugin CorrectBleach V2.0.2,” Jul. 2014.

[26] U. Weill, G. Krieger, Z. Avihou, R. Milo, M. Schuldiner, and D. Davidi, “Assessment of GFP Tag Position on Protein Localization and Growth Fitness in Yeast,” J. Mol. Biol., vol. 431, no. 3, pp. 636–641, Feb. 2019.

[27] J. M. Tkach and J. R. Glover, “Nucleocytoplasmic trafficking of the molecular chaperone Hsp104 in unstressed and heat-shocked Cells,” Traffic, vol. 9, no. 1, pp. 39–56, Jan. 2008.

[28] E. Pienaar, M. Theron, M. Nelson, and H. J. Viljoen, “A quantitative model of error accumulation during PCR amplification,” Comput. Biol. Chem., vol. 30, no. 2, pp. 102–111, Apr. 2006.

[29] L. Seppä, A.-L. Hänninen, and M. Makarow, “Upregulation of the Hsp104 chaperone at physiological temperature during recovery from thermal insult,” Mol. Microbiol., vol. 52, no. 1, pp. 217–225, Feb. 2004.

[30] Y. Sanchez and S. L. Lindquist, “HSP104 required for induced thermotolerance,” Science (80-.)., vol. 248, no. 4959, pp. 1112–1115, Jun. 1990.

[31] J. K. Heppert et al., “Comparative assessment of fluorescent proteins for in vivo imaging in an animal model system,” Mol. Biol. Cell, vol. 27, no. 22, pp. 3385–3394, Nov. 2016.

[32] I. Moore and A. Murphy, “Validating the location of fluorescent protein fusions in the endomembrane system,” Plant Cell, vol. 21, no. 6, pp. 1632–1636, Jun. 2009.

[33] K. Thorn, “Genetically encoded fluorescent tags,” Molecular Biology of the Cell, vol. 28, no. 7. American Society for Cell Biology, pp. 848–857, 01-Apr-2017.

[34] S. Teng, A. K. Srivastava, C. E. Schwartz, E. Alexov, and L. Wang, “Structural assessment of the effects of Amino Acid Substitutions on protein stability and protein-protein interaction,” Int. J. Comput. Biol. Drug Des., vol. 3, no. 4, pp. 334–349, 2010.

[35] J. Glover and R. Lum, “Remodeling of Protein Aggregates by Hsp104,” Protein Pept. Lett., vol. 16, no. 6, pp. 587–597, Jun. 2009.

[36] K. L. Schneider, T. Nyström, and P. O. Widlund, “Studying Spatial Protein Quality Control, Proteopathies, and Aging Using Different Model Misfolding Proteins in S. Cerevisiae,” Frontiers in Molecular Neuroscience, vol. 11. Frontiers Media S.A., 23-Jul-2018.

[37] C. Vacher, L. Garcia-Oroz, and D. C. Rubinsztein, “Overexpression of yeast hsp104 reduces polyglutamine aggregation and prolongs survival of a transgenic mouse model of Huntington’s disease,” Hum. Mol. Genet., vol. 14, no. 22, pp. 3425–3433, Nov. 2005.

[38] E. Balleza, J. M. Kim, and P. Cluzel, “Systematic characterization of maturation time of fluorescent proteins in living cells,” Nat. Methods, vol. 15, no. 1, pp. 47–51, Jan. 2018.

